## Supplementary Information for "Dynamic HIV-1 spike motion creates vulnerability for its membrane-bound tripod to antibody attack"

**Structure of the Env protein with three bound 4E10 Fabs**

The three 4E10 Fabs in our structure bind to the ectodomain quite differently than in the previous structures of the ectodomain in complex with three Fabs of bnAbs VRC42.01 and 10E8 (Figure S7c).^1^ The 4E10 Fabs use binding mode 2, in which the Fabs extend into the space that would normally be occupied by the membrane, which should not occur *in situ*. In contrast, in the structures of the ectodomain in complex with three VRC42.01 and 10E8 Fabs, those extend away from the membrane in the same direction as the ectodomain.

The 4E10 Fabs bind in three respects differently from each other, introducing asymmetry in the gp145•3Fab complex. First, the three Fabs are not evenly distributed around the ectodomain trimer. Rather than being separated by equal 120º angles, when viewed in the direction from the membrane to the ectodomain, the angles from Fab1 to Fab2 to Fab3 are 96º, 114º and 150º (Figure 3e). Similarly asymmetric distributions of MPER Fabs were also observed in the previously determined Env structures in complex with three VRC42.01 and 10E8 Fabs (Figure S7c). Second, the angle of the 4E10 Fab with respect to the three-fold axis of the ectodomain are 150º, 140º and 140º for Fabs 1, 2 and 3, respectively. Third, the plane of Fab1 and Fab2 is perpendicular to the membrane plane, while the plane of Fab3 is rotated by ~30º with respect to the other two Fabs. As the result of Fab1 and Fab2 being much closer together, we find that Fab1 utilizes its CDRH3 loop to also interact with the MPER-N segment from the adjacent gp145 protomer to which Fab2 is bound (Figure S7d). While Fab1 forms the interaction with its MPER epitope seen in the crystal structure, its Trp100 residue also forms aromatic stacking and hydrophobic interactions with Trp666, Leu669 and Trp670 of the neighboring MPER-N segment (Figure S7d), which is not observed for the other two Fabs.

**SUPPLEMENTARY FIGURES**

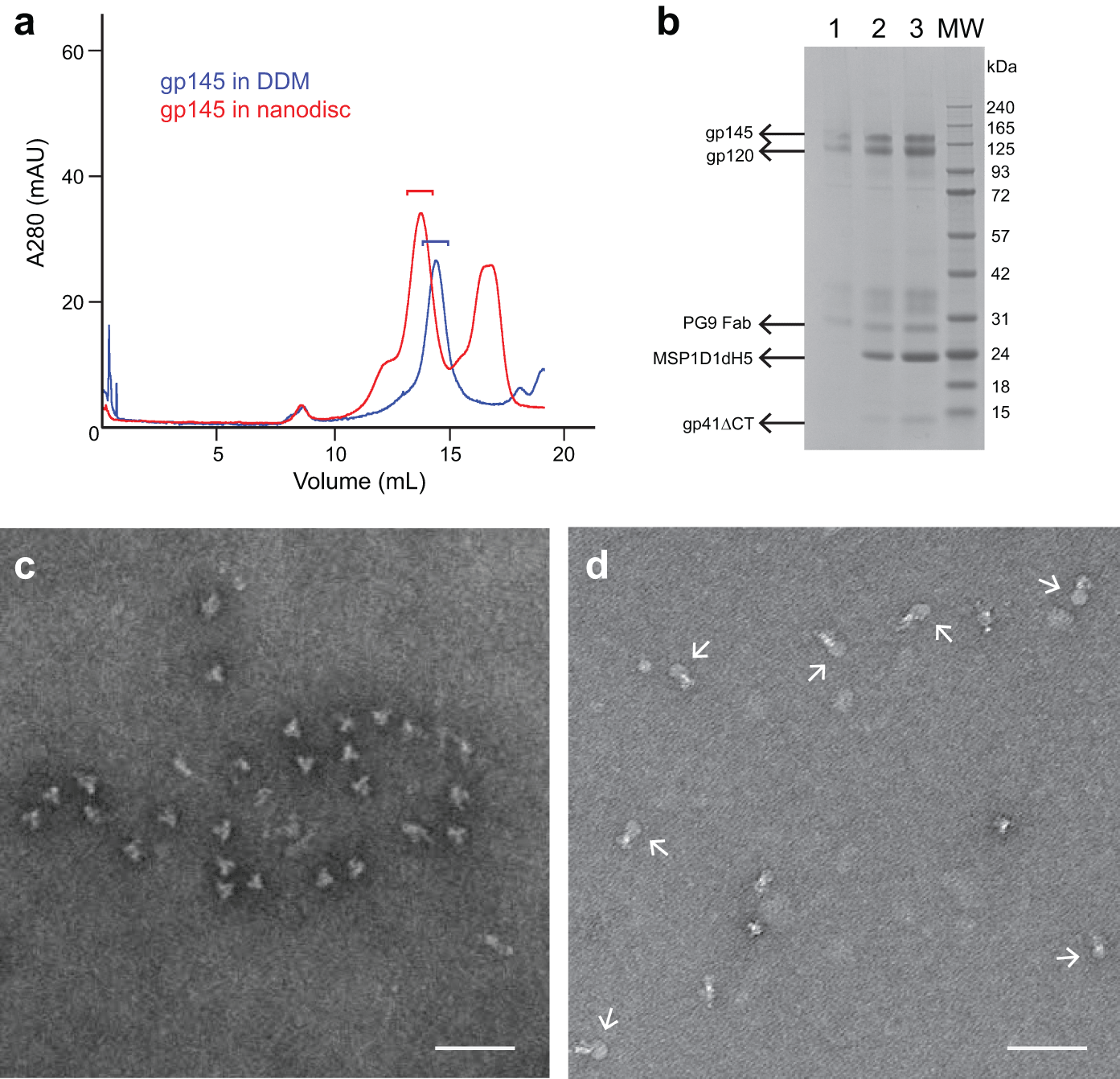

**Figure S1. Preparation of nanodisc-embedded gp145 in complex with 4E10 Fabs.** (**a**) SEC elution profiles showing a single peak for gp145 in DDM (blue line) and two peaks for gp145 in nanodiscs (red line). The peak at a lower elution volume than the peak in the DDM sample represents gp145-containing nanodiscs, whereas the peak at a higher elution volume than the peak in the DDM sample represents empty nanodiscs. Brackets indicate the fractions that were used for negative-stain EM imaging. (**b**) Coomassie Brilliant Blue-stained reducing (5% β-mercaptoethanol) SDS-PAGE gel of gp145 in DDM (lane 1) and in nanodiscs (lanes 2 and 3; sample in lane 2 is two-fold dilution of sample in lane 3). (**c** and **d**) Negative-stain EM images of gp145 in DDM (**c**) and nanodiscs (**d**). Arrows in panel (**d**) point to nanodiscs. Scale bars: 50 nm.

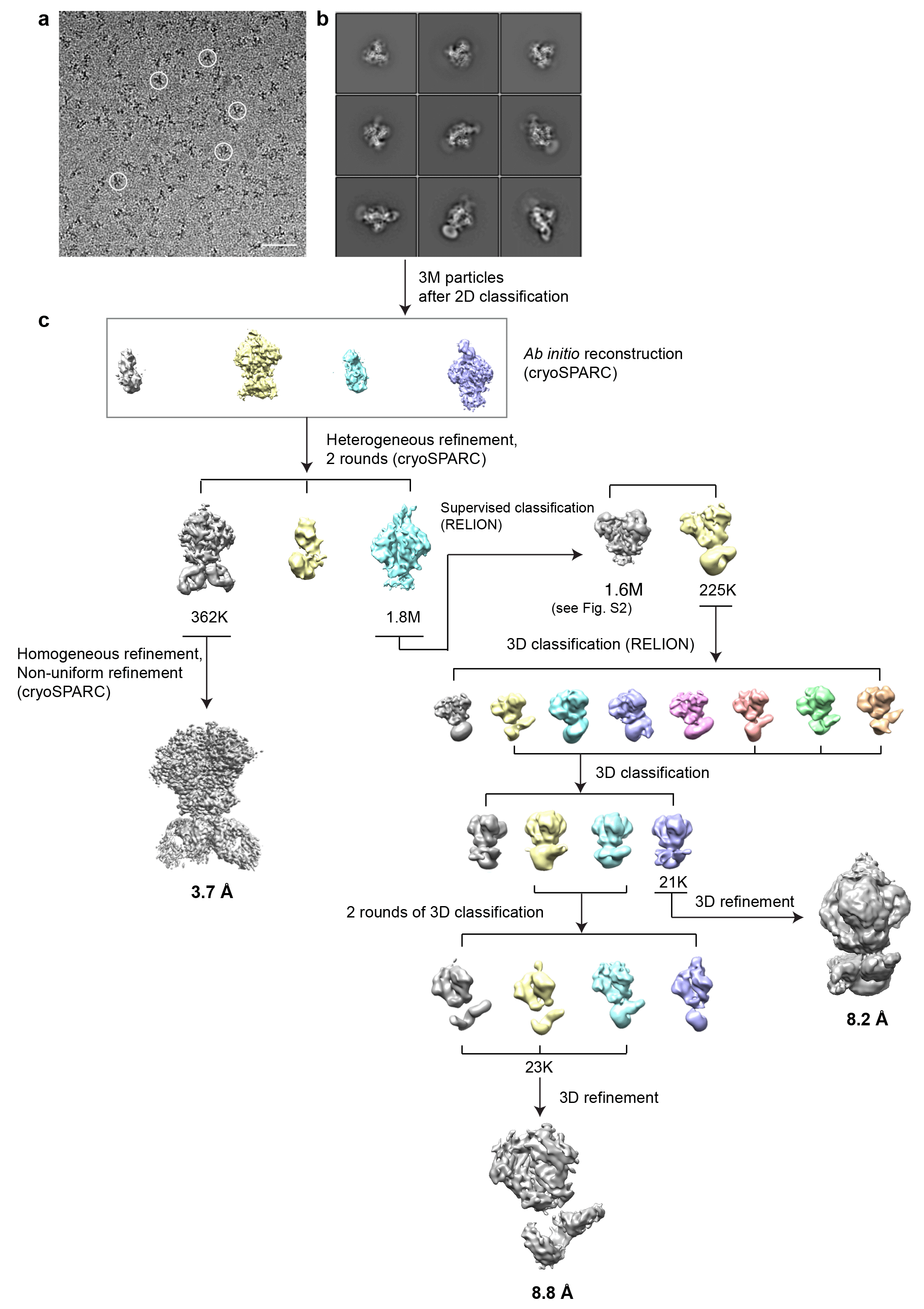

**Figure S2. Cryo-EM image processing of nanodisc-embedded gp145 incubated with 4E10 Fabs.** (**a**) Cryo-EM image of vitrified sample. Some particles are circled. Scale bar: 50 nm. (**b**) Selected 2D-class averages obtained with RELION-3. Side length of individual averages: 39.6 nm. (**c**) Image-processing workflow for 3D classification and refinement in cryoSPARC and RELION-3 that resulted in density maps of the gp145•3Fab complex at 3.7-Å resolution, the gp145•2Fab complex at 8.2-Å resolution, and the gp145•1Fab complex at 8.8-Å resolution. See Methods and Table S1 for details.

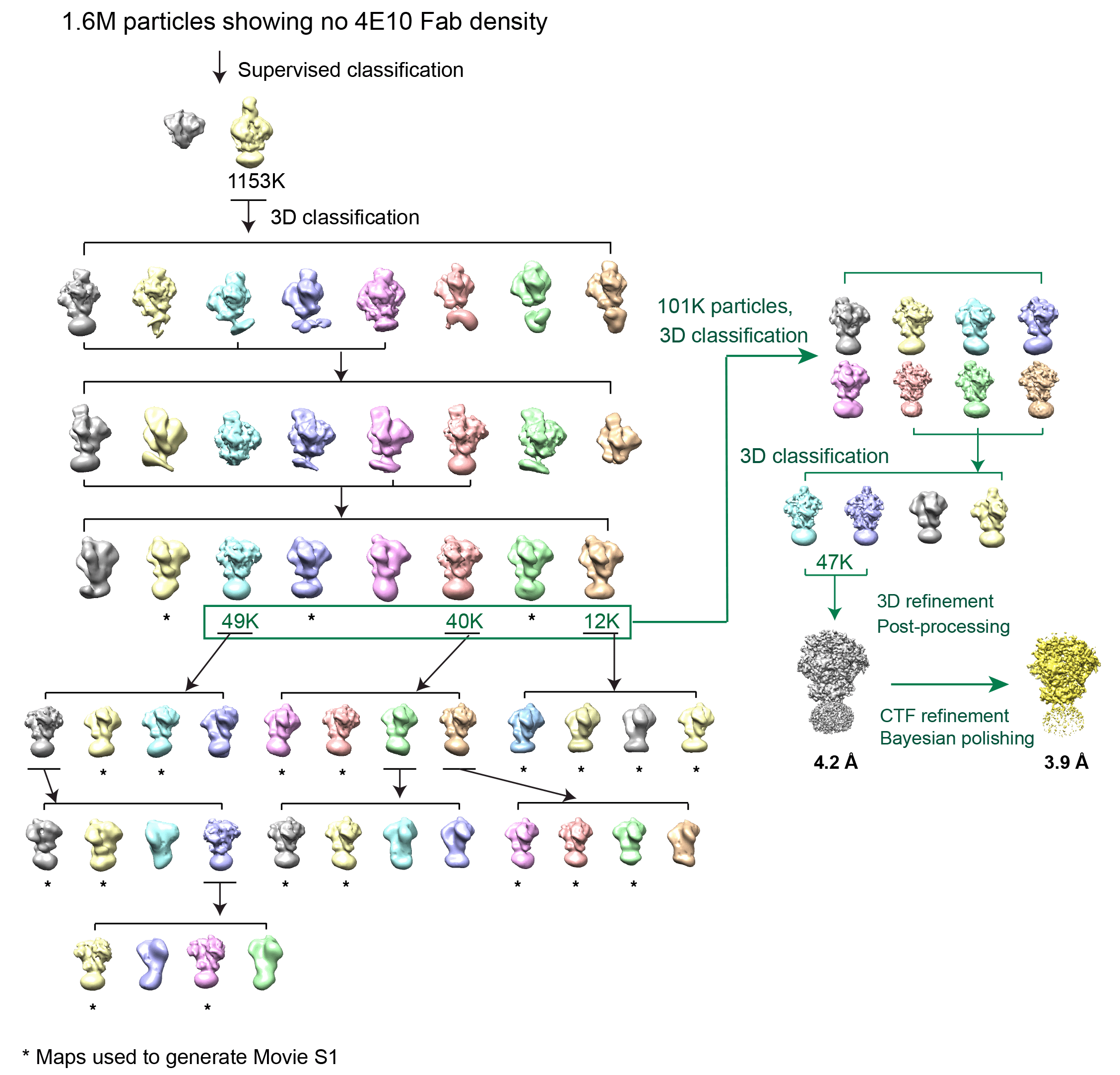

**Figure S3. Cryo-EM image processing of nanodisc-embedded gp145 incubated with 4E10 Fabs.** Continued from Figure S2. Image-processing workflow for 3D classification and refinement in RELION-3 that resulted in a density map of nanodisc-embedded gp145 without bound Fab at 3.9-Å resolution. Maps labelled with * were used to generate Movie S1. See Methods and Table S1 for details.

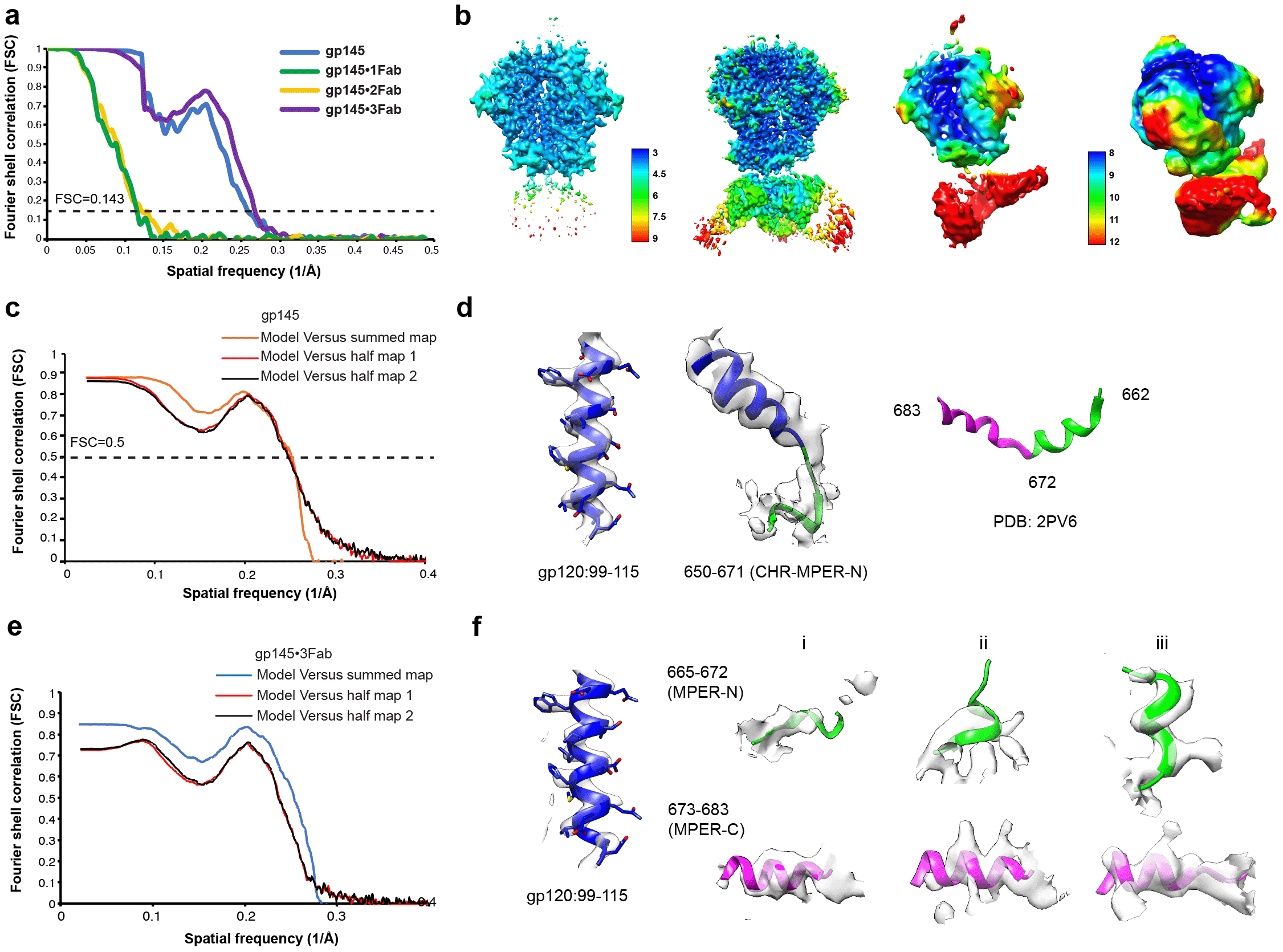

**Figure S4. FSC curves, local-resolution maps and local densities.** (**a**) FSC curves calculated between independently refined half maps for gp145 map by itself (blue), the gp145•1Fab complex (green), the gp145•2Fab complex (yellow), and the gp145•3Fab complex (purple). (**b**) From left to right: local-resolution maps as determined by the ResMap algorithm implemented in RELION-3 for gp145 by itself and the gp145•3Fab complex with the resolution scale shown in between, and local-resolution maps for the gp145•1Fab complex and the gp145•2Fab complex with the resolution scale shown in between. (**c**) Cross-validation FSC curves for nanodisc-embedded gp145 by itself: red, refined model *versus* half map 1 used for refinement (work map); black, refined model *versus* half map 2 not used for refinement (free map); orange, refined model *versus* the combined final map. The similarity of the ‘work’ and ‘free’ curves suggests no substantial over-fitting. (**d**) Selected local cryo-EM densities for the map of gp145 by itself (left) and the NMR structure of the MPER used for docking of the N-segment into the cryo-EM density. (**e**) Cross-validation FSC curves for nanodisc-embedded gp145•3Fab complex: red, refined model *versus* half map 1 used for refinement (work map); black, refined model *versus* half map 2 not used for refinement (free map); orange, refined model *versus* the combined final map. The similarity of the ‘work’ and ‘free’ curves suggests no substantial over-fitting. (**f**) Selected local cryo-EM densities for the map of the gp145•3Fab complex. The densities representing the MPER-N and MPER-C segments bound by Fab1 (i), Fab2 (ii) and Fab3 (iii) are shown.

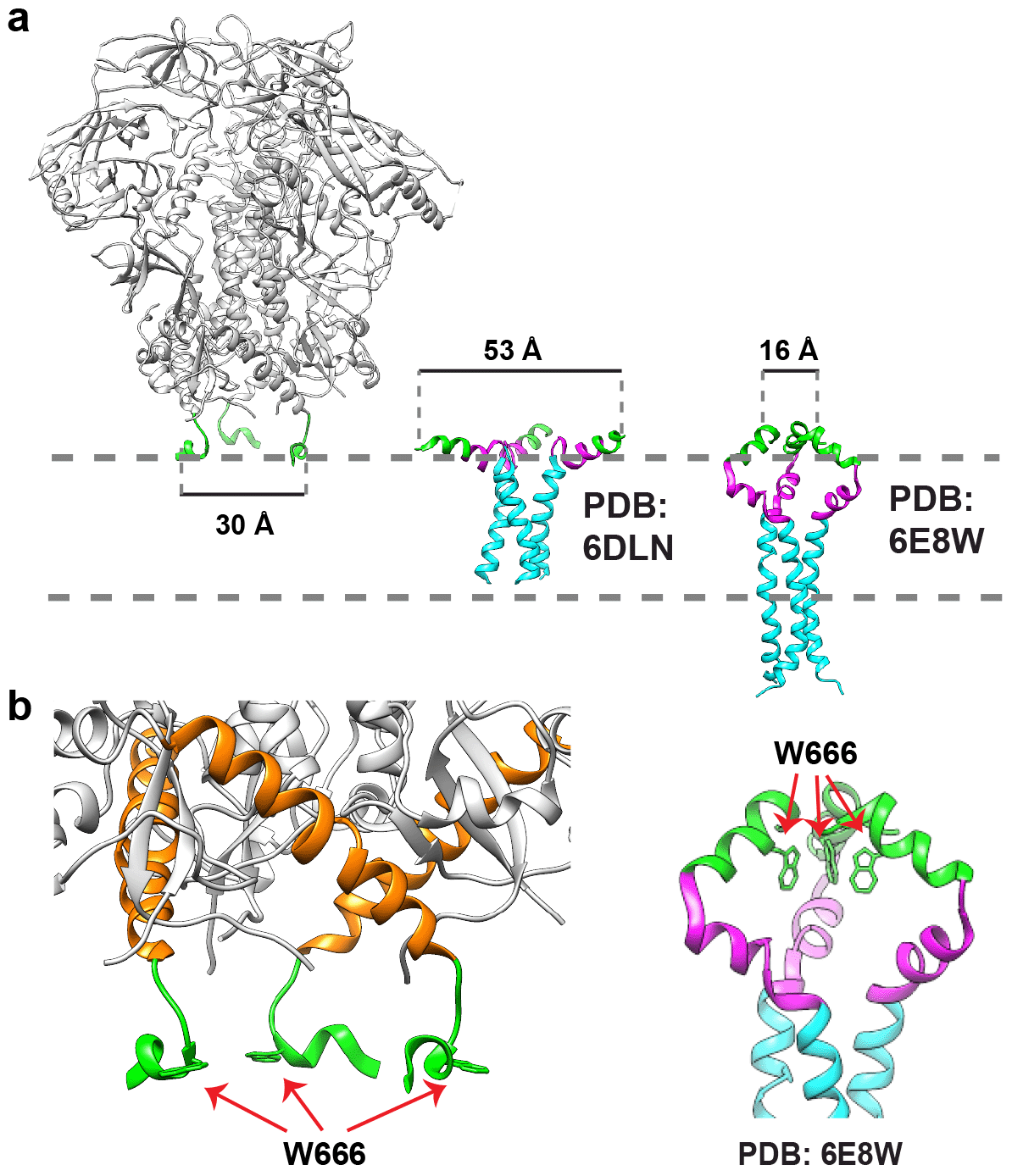

**Figure S5. Comparison of the MPER segments in the cryo-EM structure of nanodisc-embedded gp145 with those in NMR structures.** (**a**) The distance between the N-terminal ends of the MPER segments, measured between the Cα atoms of residue Lys665 from adjacent protomers, in the cryo-EM structure (this study) and two previously determined NMR structures (PDB: 6DLN and PDB: 6E8W).^2,3^ Dashed grey lines indicate the membrane. (**b**) Comparison of the dispersed MPER-N segments in the cryo-EM structure (this study; left panel) with the converged MPER-N segments in one of the NMR structures (PDB: 6E8W; right panel).^3^ The side chains of residue Trp666 are shown in stick representation.

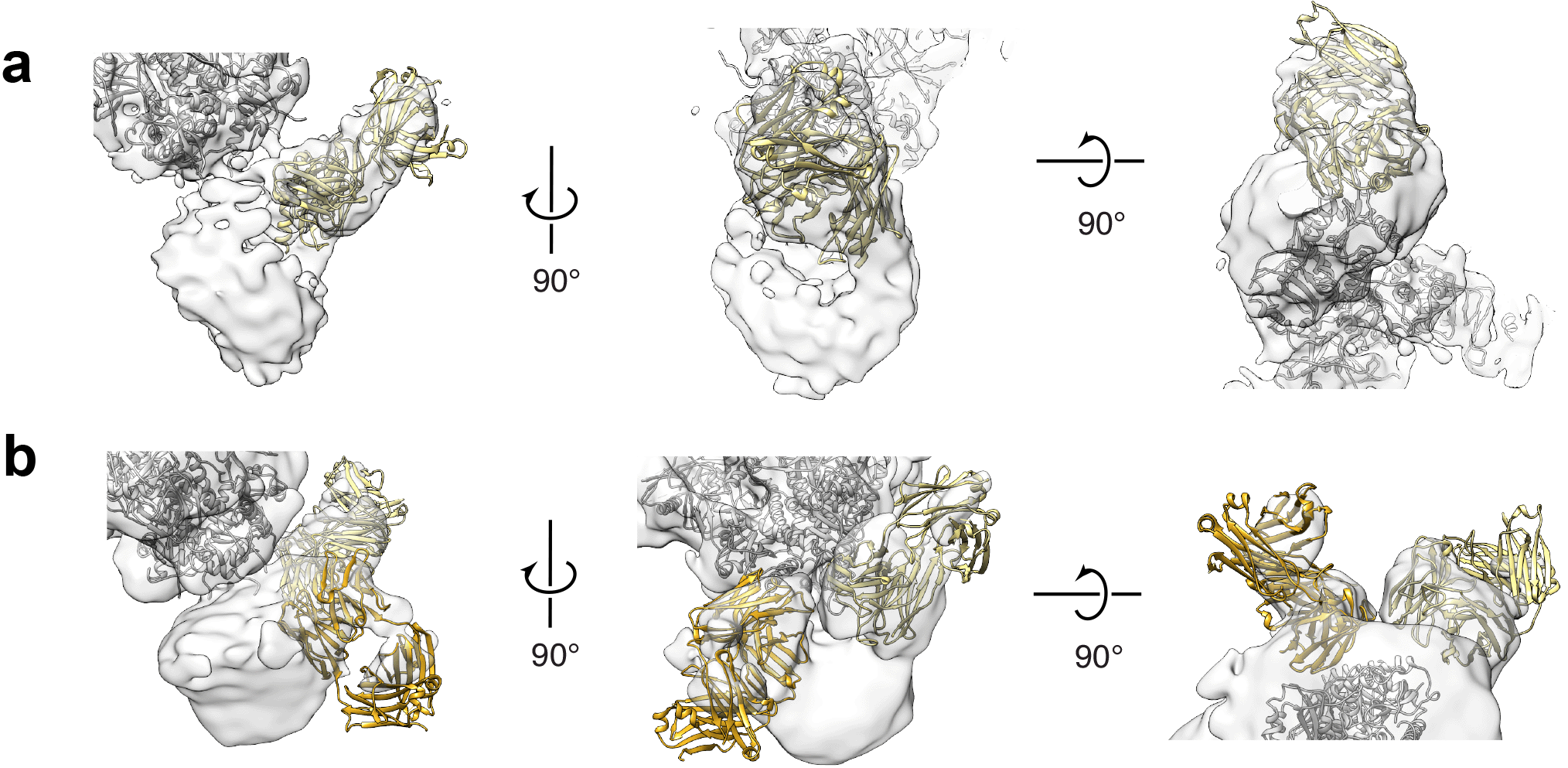

**Figure S6. Docking of the 4E10 Fab crystal structure into the maps of nanodisc-embedded gp145 in complex with one Fab (gp145•1Fab) and two Fabs (gp145•2Fab).** (**a** and **b**) Three orthogonal views that show the docking of the 4E10 Fab crystal structure (PDB: 4XC3)^4^ into the gp145•1Fab map (**a**) and the gp145•2Fab map (**b**).

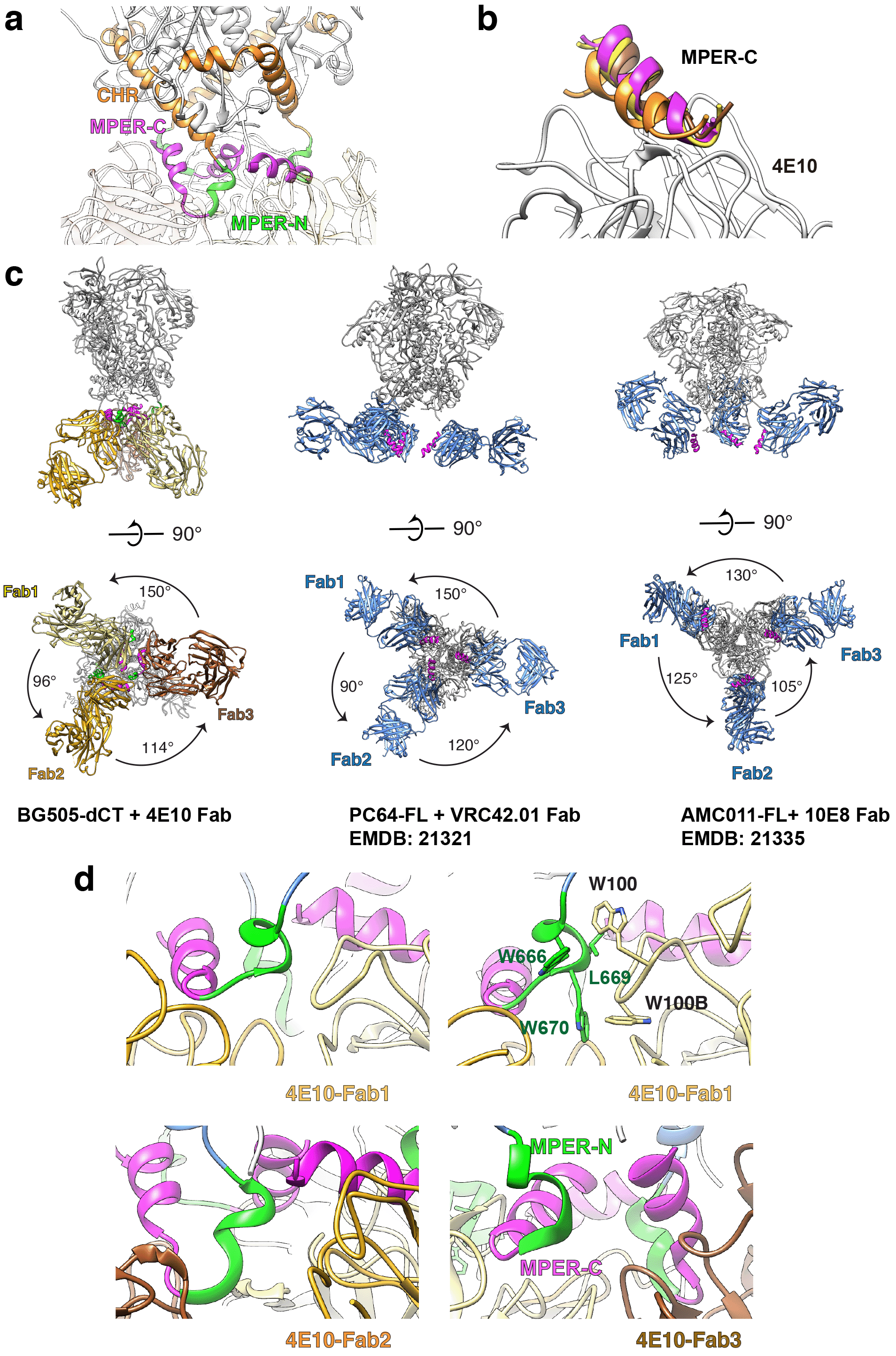

**Figure S7. Connection of the MPERs to the ectodomain and their interaction with bnAbs.** (**a**) Close-up view of the connections of the MPERs with the CHRs of the ectodomain in the gp145•3Fab structure. (**b**) Interaction of the three MPER-C segments in the gp145•3Fab structure (yellow, gold and brown) and the MPER-C epitope in the crystal structure (magenta) with the 4E10 Fab. The structures were overlaid based on the Fab structure. (**c**) Cryo-EM structures of HIV-1 Env in complex with three anti-MPER bnAb Fabs seen parallel (top) and perpendicular to the membrane plane (bottom). Left panels: BG505-dCT gp145 with three 4E10 Fabs (this study). All three Fabs are bound in mode 2 and extend into the space that would normally be occupied by the membrane. Middle panels: Map: PC64 gp160 with three VRC42.01 Fabs (EMDB:21321)^1^; the coordinates used for docking are 5I8H for the gp160 ectodomain^5^ and 6MTP for the VRC42.04 Fab in complex with the gp41 peptide.^6^ Right panels: AMC011 gp160 with three 10E8 Fabs (PDB: 6PVX).^1^ The Fabs seen in the two previous structures (middle and right panels) are bound to Env in mode 1 and extend away from the membrane. The angles between the three Fabs differ in the three structures. (**d**) The region between the 4E10 Fabs and the MPER-N segments from the neighboring protomers. Fab1 is closer to the neighboring MPER-N segment, possibly forming hydrophobic interactions. The residues involved in these putative interactions are shown in stick representation. Fabs 2 and 3 are further away from their neighboring MPER-N segments and are thus unlikely to from hydrophobic interactions. Green and magenta colors are used to delineate the MPER-N and MPER-C segments, respectively.

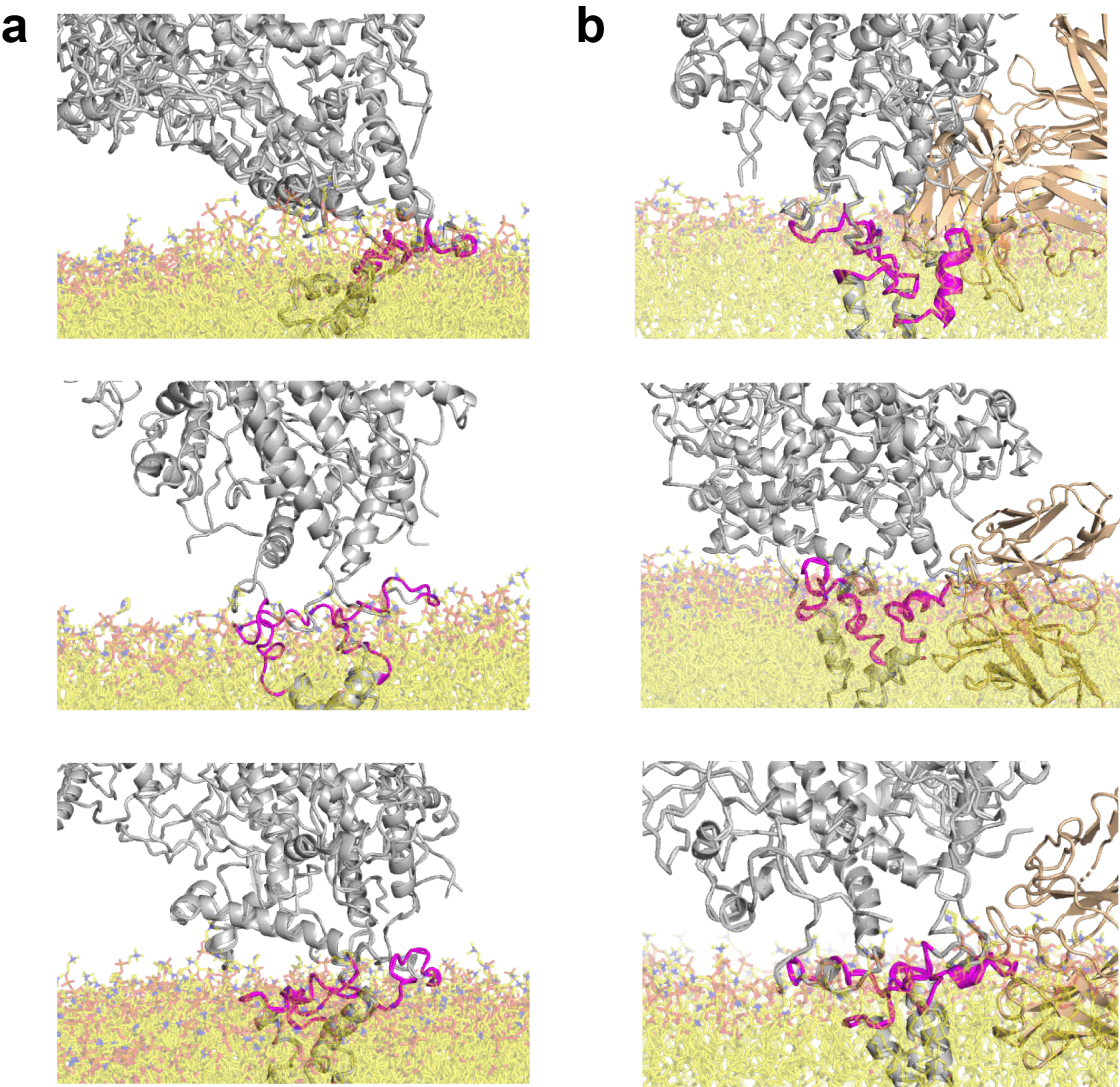

**Figure S8. All-atom views generated from snapshots of the coarse-grained molecular-dynamics simulations of membrane-embedded gp145 showing the ectodomain at high tilts.** (**a**) Three snapshots in which the MPER-C segment lost its α-helical secondary structure, thus preventing reliable docking of the 4E10 Fab. (**b**) Three snapshots in which the MPER-C segment remained α-helical, but where docking of the 4E10 Fab resulted in steric clashes with the ectodomain or the Fab being partially buried in the lipid bilayer.

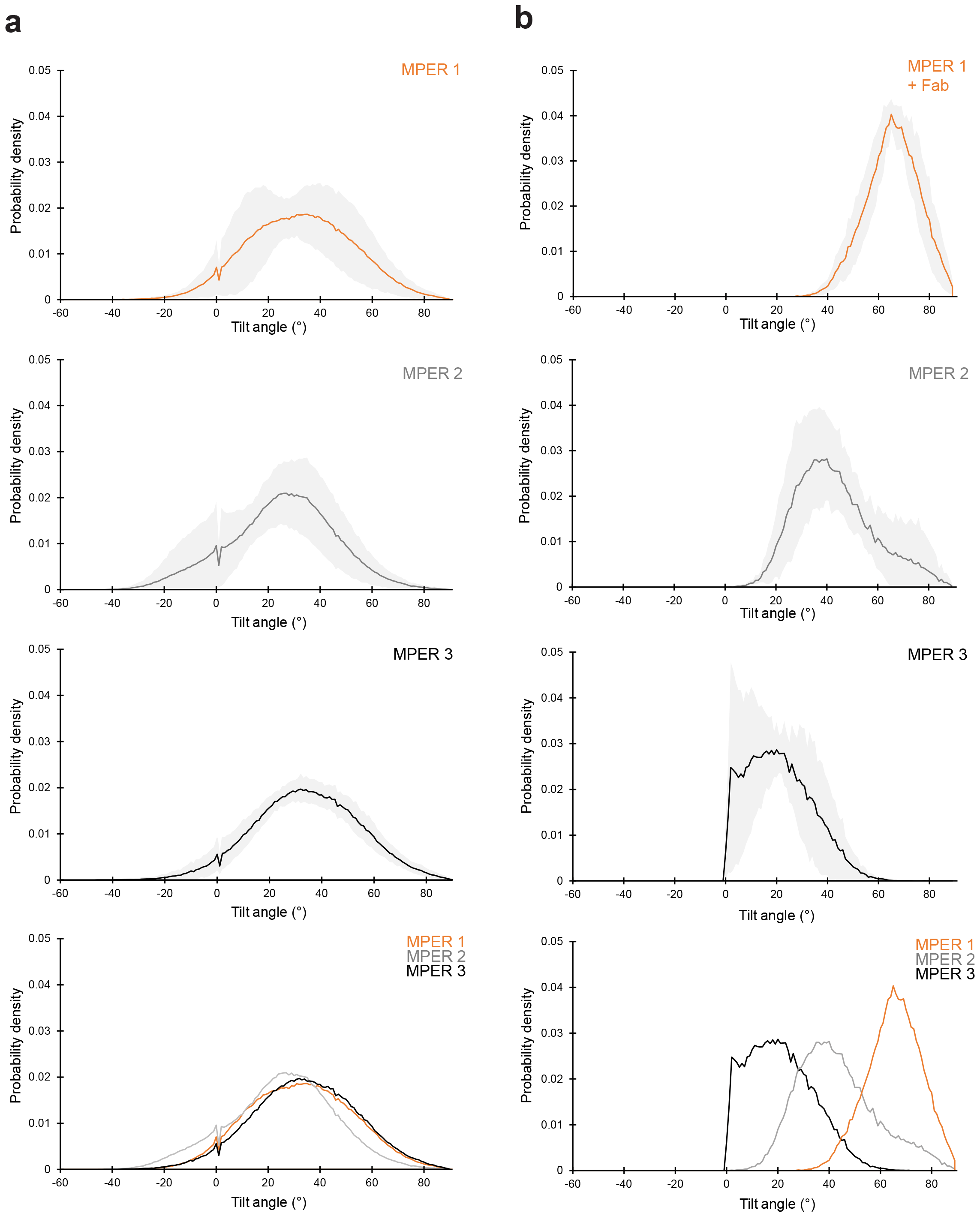

**Figure S9. Distribution of the angles adopted by the three MPER-C segments in coarse-grained molecular-dynamics simulations of membrane-embedded gp145.** (**a**) Graphs showing the distribution of the angles adopted by the MPER-C segments in unliganded gp145, summarizing data from all five repeats. The top three panels show the angle distribution for the three MPER-C segments individually, with the solid lines representing the average value and the grey bands indicating the standard deviation. The bottom panel shows an overlay of the angle distribution for the three MPER-C segments, demonstrating that the three MPER-C segments in unliganded gp145 show comparable behavior. (**b**) Graphs showing the distribution of the angles adopted by the MPER-C segments in 4E10 Fab-bound gp145, summarizing data from all five repeats. The top three panels show the angle distribution for the three MPER-C segments individually, with the solid lines representing the average value and the grey bands indicating the standard deviation. The bottom panel shows an overlay of the angle distribution for the three MPER-C segments, demonstrating that the three MPER-C segments in 4E10 Fab-bound gp145 adopt distinct angles.

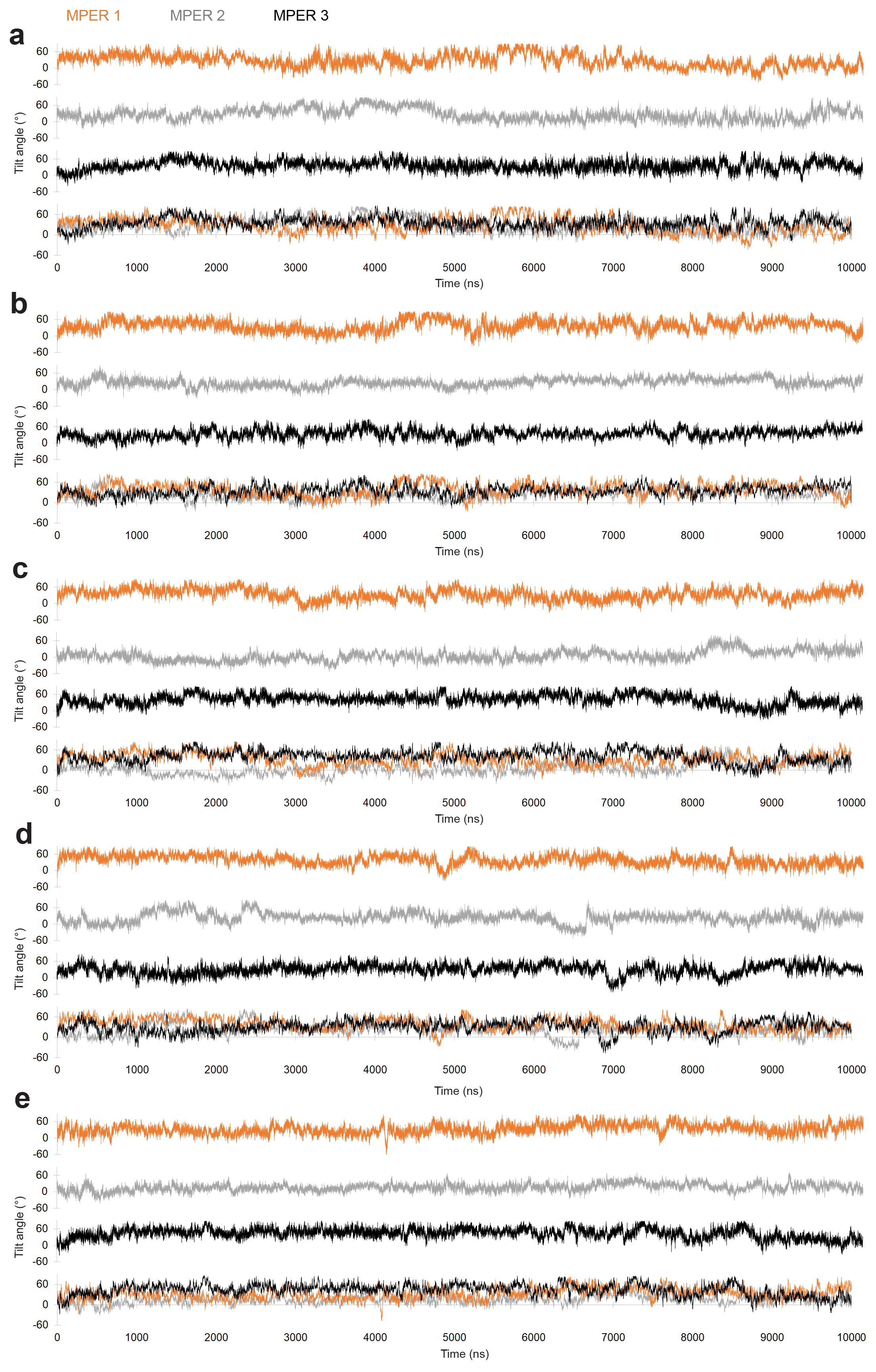

**Figure S10. Angles of the three MPER-C segments adopted over time in coarse-grained molecular-dynamics simulations of unliganded gp145.** (**a**-**e**) For each of the five repeats, the angles adopted by each MPER-C segment is shown on their own (top three panels) as well as overlaid (bottom panel).

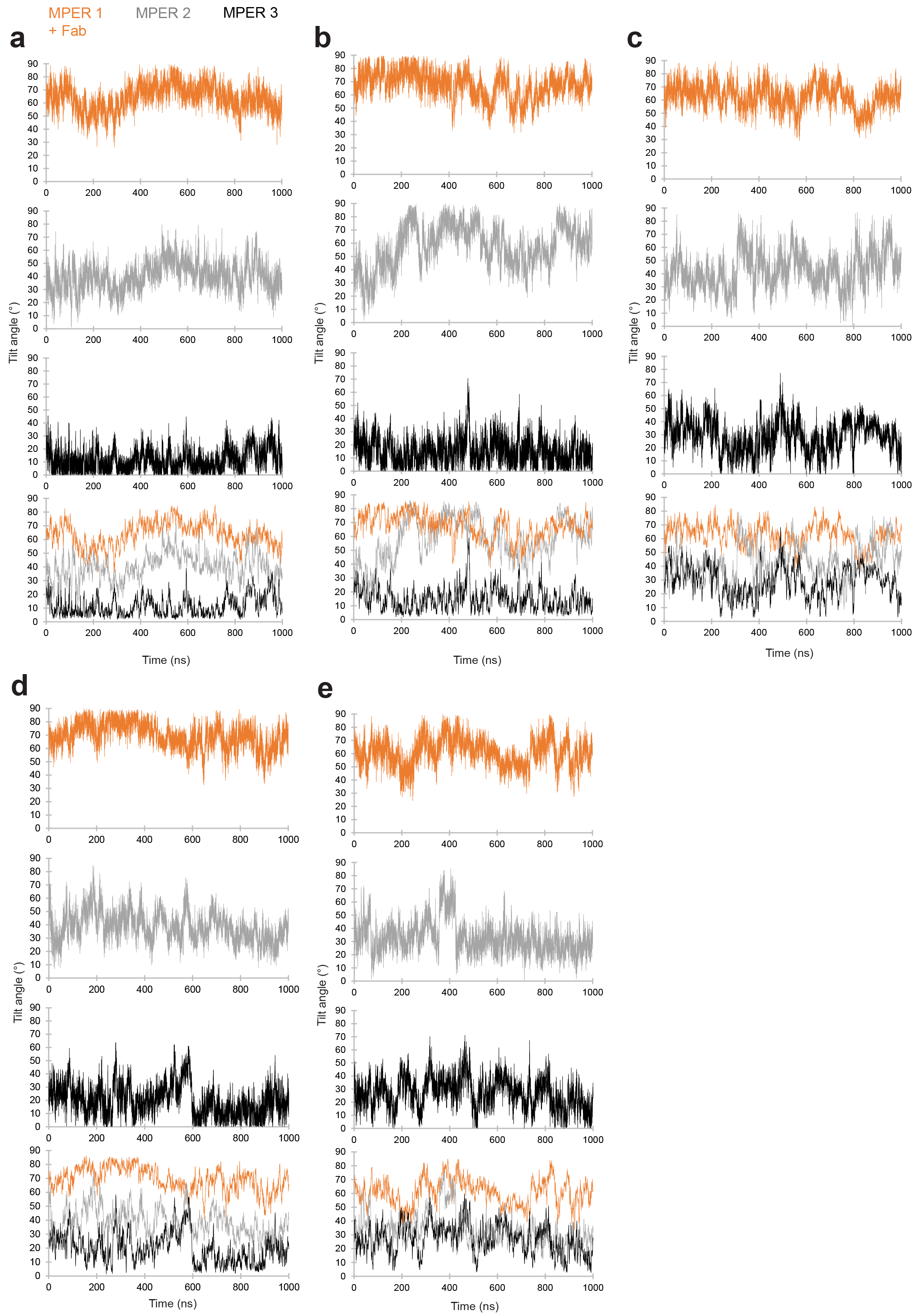

**Figure S11. Angles of the three MPER-C segments adopted over time in coarse-grained molecular-dynamics simulations of 4E10 Fab-bound gp145.** (**a**-**e**) For each of the five repeats, the angles adopted by each MPER-C segment is shown on their own (top three panels) as well as overlaid (bottom panel).

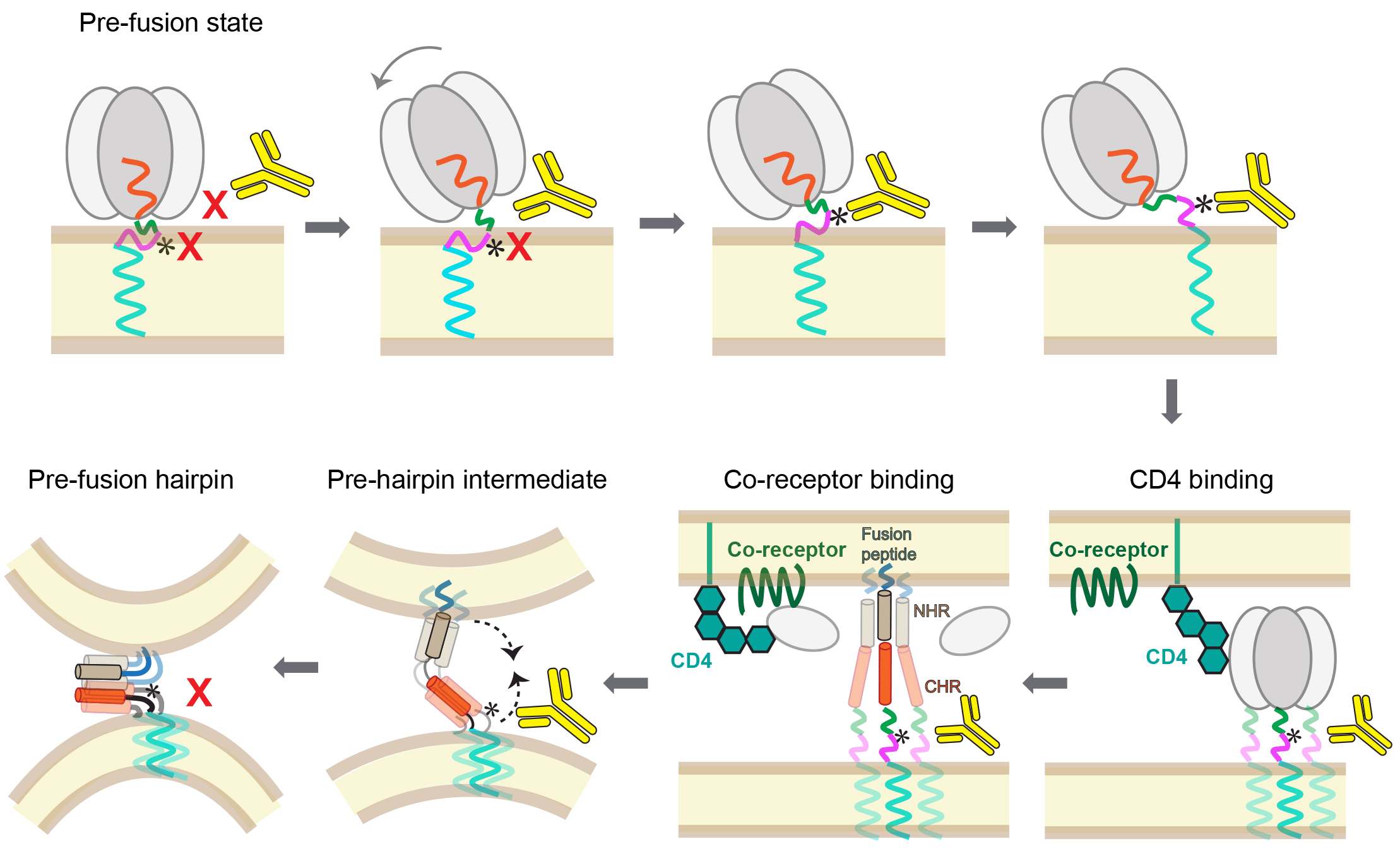

**Figure S12. The stepwise binding of the anti-MPER bnAb 4E10 to the Env protein and exposure of MPER epitope during different states of the fusion process.** Upper panels, see Figure 6a. Lower panels: While the current study establishes that bnAb 4E10 (and possibly other anti-MPER bnAbs) can bind to HIV-1 Env in the prefusion state, the epitope (*) may also be accessible after CD4 engagement, gp120 shedding and during the pre-hairpin intermediate state. Once the pre-fusion hairpin has formed, however, the epitope is likely obscured. For simplicity of illustration the conjoint CD4 and chemokine receptor binding to the gp120 protomer in the unshed state is omitted.

**Table S1. Cryo-EM data collection, refinement and validation statistics**

|  | **gp145**  **(EMDB-25022)**  **(PDB: 7SC5)** | **gp145+ 3•4E10 Fab**  **(EMDB-25045)**  **(PDB: 7SD3)** | **gp145+ 2•4E10 Fab**  **(EMDB-25025)** | **gp145+ 1•4E10 Fab**  **(EMDB-25024)** |
| --- | --- | --- | --- | --- |
| **Data collection and processing** |  | | | |
| Microscope | Titan Krios | | | |
| Voltage (kV) | 300 | | | |
| Detector | K2 Summit (Gatan) | | | |
| Pixel size (Å) | 1.03 | | | |
| Magnification | 29,000 | | | |
| Defocus range (µm) | -1.5 to -3.0 | | | |
| Total dose (electrons/Å^2^) | 80 | | | |
| Movie stacks(no.) | 30404 | | | |
| Initial particle images(no.) | 3,039,896 | | | |
| Particle images for final reconstruction (no.) | 47,616 | 362,646 | 21,454 | 23,583 |
| Symmetry imposed | C3 | C1 | C1 | C1 |
| Map resolution at 0.143 FSC threshold (Å) | 3.9 | 3.7 | 8.2 | 8.8 |
| **Refinement** | | |  |  |
| Initial models used (PDB code) | 5I8H | 5I8H; 4XC3 |  |  |
| Map sharpening B factor (Å^2^) | -110 | -120 |  |  |
| Model composition |  |  |  |  |
| Non-hydrogen atoms | 14703 | 25256 |  |  |
| Protein residues | 1770 | 3111 |  |  |
| Ligands | BMA:6 NAG:54 | BMA:6 NAG:54 |  |  |
| B factors (Å^2^) |  |  |  |  |
| Protein | 130.15 | 134.33 |  |  |
| Ligand | 158.76 | 134.86 |  |  |
| R.m.s. deviations | | |  |  |
| Bond lengths (Å) | 0.008 | 0.011 |  |  |
| Bond angles (°) | 1.248 | 1.475 |  |  |
| MolProbity score | 2.11 | 2.77 |  |  |
| Clashscore | 10.05 | 11.23 |  |  |
| Poor rotamers (%) | 0.96 | 7.23 |  |  |
| Ramachandran plot (%) | | |  |  |
| Favored/Allowed/Outliers | 88.47/11.3/0.23 | 90.08/9.86/0.07 |  |  |

**Movie S1. Motions of the HIV-1 Env ectodomain on the membrane surface**. The movie was generated by morphing between 20 ectodomain orientations obtained by docking the ectodomain structure into EM maps obtained during image processing (Fig. S3).

**Movie S2. Largest tilts of the HIV-1 Env ectodomain relative to the membrane surface**. The movie was generated by morphing between the two ectodomain orientations showing the highest tilt angles.

**Movie S3. Coarse-grained molecular-dynamics simulation of membrane-embedded gp145.** The movie shows 2 μs of one of the repeats. Note the spontaneous tilting of the ectodomain and the uncorrelated dynamics of the three MPER segments (MPER-N in green and MPER-C in magenta).

**Movie S4. Coarse-grained molecular-dynamics simulation of membrane-embedded gp145 with bound 4E10 Fab.** The movie shows 0.9 μs of one of the repeats. Note that the ectodomain still adopts different tilts but that the average tilt is higher than in the unliganded gp145. Fab binding appears to stabilize the three MPER segments at different average angles (MPER-N in green and MPER-C in magenta).
